# Quieting the Storm: Hypoxia as a Strategy to Boost UC-MSC Therapies for Neonatal Hypoxic-Ischemic Encephalopathy

**DOI:** 10.1101/2025.01.21.634045

**Authors:** Inês Serrenho, Vera Mendes, Inês Caramelo, Carla M. Cardoso, Bruno Manadas, Graça Baltazar

## Abstract

Integrating stem cell therapies into clinical settings faces several challenges, particularly in achieving the high cell yields necessary for attaining therapeutic doses. Preconditioning with hypoxic conditions has shown promise in enhancing the UC-MSCs reparative capabilities of the central nervous system. Recent evidence suggests that oxygen concentration and exposure duration can shape MSCs’ phenotypes, supporting the need for further optimization of this strategy in a way to achieve maximal repair. In this study, we assessed the effects of both prolonged mild hypoxia (MH; 5% oxygen for 48 hours) and short severe hypoxia (SSH; 0.1% oxygen for 24 hours) on UC-MSCs’ ability to alleviate motor and cognitive deficits in a rodent model of neonatal HIE. Our results show that short, severe hypoxia led to more improvements in functional recovery than prolonged mild hypoxia, supporting that specific preconditioning parameters are crucial in maximizing UC-MSC therapeutic potential. To investigate the molecular effects of hypoxia-preconditioned MSCs in the neonatal brain post-HIE, we employed untargeted proteomics on ipsilesional brain samples from control, HIE, HIE treated with naïve UC-MSCs, and HIE treated with SSH-preconditioned UC-MSCs groups, 30 days after lesion induction. This approach identified protein signatures related to injury and therapeutic intervention. Pathway enrichment analysis further revealed that administration of UC-MSCs preconditioned with short severe hypoxia significantly impacted neural signaling, protein synthesis, and energy metabolism pathways, pointing to long-term mechanisms that may support neuronal repair. These findings enhance our understanding of hypoxia-preconditioning in MSCs therapy in driving a positive therapeutic response, supporting the development of more effective and feasible treatments for neonatal HIE.

## 1. Introduction

MSCs have become a promising strategy for regenerative medicine (1). These cells can promote tissue repair and modulate disrupted processes after brain injury (2, 3). MSCs promote neurorepair through various mechanisms, including the secretion of growth factors, reduction of neuroinflammation, including modulation of glial response, and stimulation of neurogenesis and angiogenesis (1, 4, 5). Although still under clinical and preclinical investigation, MSC-based therapies have opened new avenues for treating neonatal HIE by targeting the underlying neuroinflammatory and neurodegenerative processes (4, 6, 7). In models of HIE, MSCs have the potential to mitigate the secondary injury cascade by modulating inflammatory response, stabilizing the blood-brain barrier, and reducing neuronal damage, which together lead to improved cognitive and motor outcomes (7, 8). However, MSCs-based therapy faces several challenges that limit its clinical applicability. One of the primary caveats is the need for high cell yields to achieve therapeutic efficacy, which can be challenging to produce and maintain under clinical-grade conditions (9, 10). Preconditioning MSCs with hypoxia has emerged as a promising strategy to enhance their therapeutic potential (11–22). Hypoxic preconditioning can activate adaptive cellular responses in MSCs, making them more resilient and better prepared to survive and function in the post-injury environment. This strategy also improves MSC regenerative capacity by upregulating factors that support cell migration, engraftment, and paracrine signaling, which are essential for neurorepair (11, 13–16, 21, 23–30).

Recent evidence suggests that hypoxic preconditioning could harness differential effects on MSCs depending on the oxygen pressure used and the duration of the stimulus (31). Indeed, the reported heterogenous results concerning the impact of hypoxia on MSCs’ therapeutic capacity could be explained by the variations in the oxygen pressure used to precondition the cells that usually vary between 0-7% [reviewed by Samal et al., 2021 (31)]. The authors suggest that most hypoxic preconditioning protocols fall within one of three different categories: prolonged severe exposure (<0.1% oxygen, more than 24 hours), prolonged mild exposure (2-5% oxygen, more than 24 hours), short severe exposure (<0.1% oxygen, less than 24 hours) and short mild exposure (2-5% oxygen, less than 24 hours) (31). Prolonged exposure to severe hypoxia is associated with reduced MSC proliferation and increased apoptosis, whereas longer exposure to mild hypoxia enhances chondrogenesis and proliferation (32–36). These reports highlight the need to identify which protocols of hypoxia preconditioning can maximize the therapeutic potential of MSCs and their ability to repair brain injuries.

The main aim of this study was to evaluate how two hypoxic preconditioning protocols— prolonged mild hypoxia (MH; 5% oxygen for 48 hours) and short severe hypoxia (SSH; 0.1% oxygen for 24 hours)—could influence the therapeutic efficacy of UC-MSCs in mitigating deficits associated with neonatal HIE and how its administration promotes brain recovery in chronic phases of this condition.

## 2. Results

### 2.1. UC-MSCs exposed to different protocols of hypoxia preconditioning elicit distinct functional recovery in HIE rats

To assess the effects of different preconditioning strategies on neurobehavioral outcomes commonly affected by neonatal HIE, we employed the Rice-Vannucci model to induce an HI brain lesion in P10 rats. Two days post-injury, animals received an IV administration of a sub-optimal dose of UC-MSCs (50,000 cells), selected based on prior optimization (Supplementary Figure 1). Before administration, cells were subjected to different oxygen preconditioning treatments. The experimental groups used were:

- Control: unlesioned animals, IV administration of PBS.
- HIE: HI brain lesion, IV administration of PBS.
- HIE+N-MSC: HI brain lesion, IV administration of UC-MSCs exposed to 21% oxygen for 24 hours (naïve UC-MSCs).
- HIE+MH-MSC: HI brain lesion, IV administration of UC-MSCs exposed to 5% oxygen for 48 hours (MH preconditioning).
- HIE+SSH-MSC: HI brain lesion, IV administration of UC-MSCs exposed to 0.1% oxygen for 24 hours (SSH preconditioning).

The negative geotaxis reflex test was used to evaluate vestibular function and motor development in neonatal rats at P14 and P17 (Figure 1A). At P14, control animals completed the task in 1.43 ± 0.15 seconds, whereas HI-injured animals required three times more time (4.68 ± 0.56 seconds, p < 0.0001 compared to control). Treatment with SSH-preconditioned UC-MSCs (HIE+SSH-MSC group) resulted in marked improvement in the performance (2.15 ± 0.50 seconds, p = 0.0003 compared to HIE). Animals from HIE+MH-MSC and HIE+N-MSC groups demonstrated no significant improvements (3.24 ± 0.35 and 3.77 ± 0.18 seconds, respectively). The therapeutic effects of MSCs administration on this parameter became more pronounced at P17. While control animals maintained consistent performance (1.02 ± 0.18 seconds), HIE animals maintained the impairment (3.32 ± 0.47 seconds, p = 0.0013 compared to control). The HIE+SSH-MSC group demonstrated the most robust recovery (1.04 ± 0.25 seconds, p = 0.0022 compared to HIE), significantly outperforming both HIE+N-MSC and HIE+MH-MSC groups (p < 0.0001 and p = 0.0031, compared to HIE+SSH-MSC, respectively).

**Figure 1.**
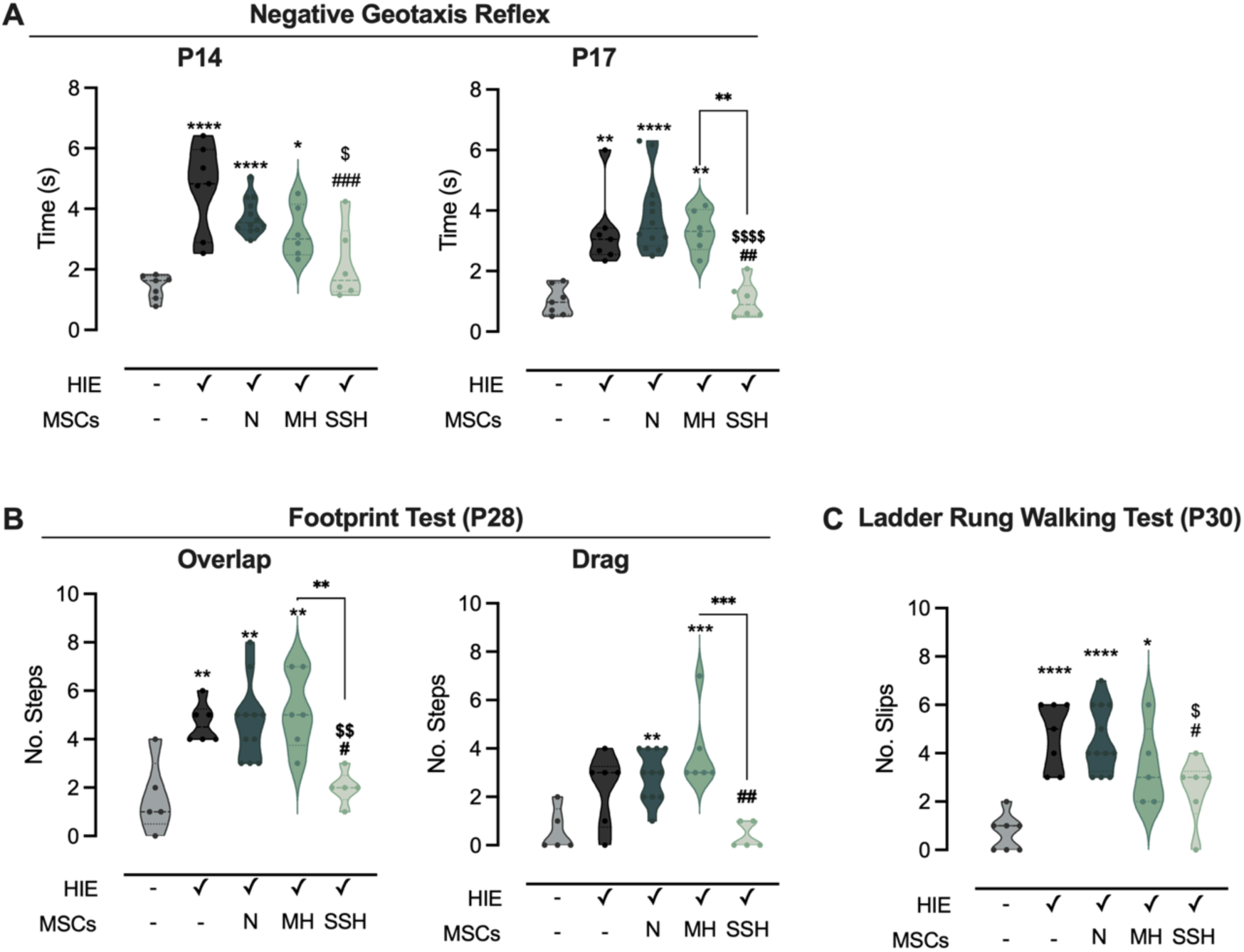
Impact of N-, SSH-or MH-UC-MSCs administration on the sensorimotor functions in HI-injured rats. (A) Latency (seconds) for pups to rotate 180° and face uphill in the negative geotaxis reflex test at P14 and P17. (B) Footprint analysis assessing dragging and foot overlap at P28. (C) Number of slips in the ladder rung walking test at P30. Statistical analysis was performed using one-way ANOVA coupled with Tukey’s multiple comparison test for the negative geotaxis reflex test and ladder rung walking test; and the Kruskal–Wallis test and Dunn’s multiple comparison test for the footprint test. Statistical differences are indicated as: * p < 0.05, ** p < 0.01, and **** p < 0.0001 vs Control (except when the comparison is identified with brackets); ^#^ p < 0.05, ^##^ p < 0.01, and ^###^ p < 0.001 vs HIE; ^$^ p < 0.05, ^$$^ p < 0.01, and ^$$$$^ p < 0.0001 vs HIE+N-MSC.

Regarding the impact of injury and treatment on the stride patterns (Figure 1B), HIE animals exhibited a tendency to have a higher number of dragging steps (2 ± 0.6 steps) compared to controls (1 ± 0.4 steps). HIE+N-MSC and HIE+MH-MSC groups still presented stride impairments when compared to the control animals (3 ± 0.3 steps, p = 0.0097; and 4 ± 0.7 steps, p = 0.001, respectively).

Analysis of step overlapping also revealed impairment in HIE animals (5 ± 0.3 steps) compared to controls (2 ± 0.7 steps, p = 0.0084). HIE+SSH-MSC treatment led to substantial improvement (2 ± 0.3 steps, p = 0.0269 vs. HIE), outperforming both HIE+N-MSC (5 ± 0.5 steps, p = 0.0085 vs. HIE+SSH-MSC) and HIE+MH-MSC (5 ± 0.7 steps, p = 0.0062 vs. HIE+SSH-MSC) groups.

Motor coordination and limb placement accuracy were affected by the HIE lesion, as showed by the increase in the number of slips (5 ± 0.5 slips, Figure 1C) compared to controls (1 ± 0.3 slips, p < 0.0001). This impairment was reversed in the HIE+SSH-MSC group (3 ± 0.6 slips, p = 0.0391 vs. HIE). Moreover, animals in this group outperformed HIE+N-MSC animals in this test (5 ± 0.4 slips, p = 0.0280 vs. HIE+SSH-MSC), although being no different from HIE+MH-MSC group (3 ± 0.7 slips).

Cognitive function, in specific recognition memory, was assessed using the novel object recognition test at two different developmental stages -P21, corresponding to human infancy, and P38, corresponding to early adulthood (Figure 2).

**Figure 2.**
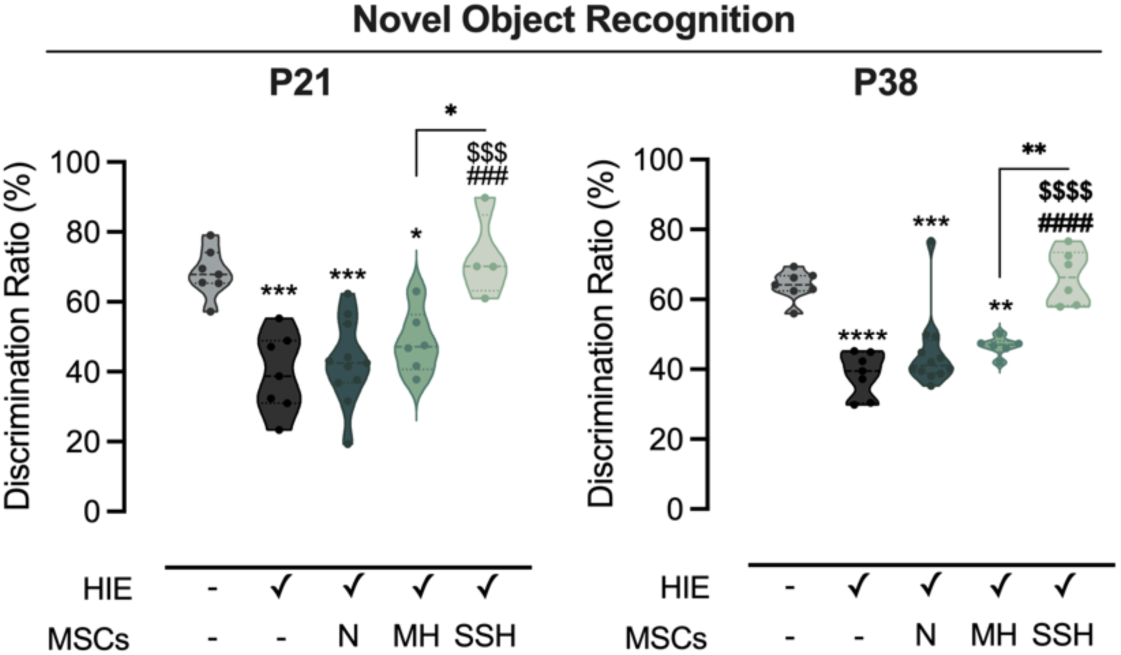
Impact of N-, SSH-or MH-UC-MSCs administration on recognition memory in HI-injured rats. Discrimination ratio reflecting a preference for the novel object during the testing phase of the novel object recognition test at P21 and P38. Statistical analysis was performed using one-way ANOVA followed by Tukey’s multiple comparison test. Statistical differences are indicated as: * p < 0.05, ** p < 0.01, *** p < 0.001, and **** p < 0.0001 vs Control (except when the comparison is identified with brackets); ### p < 0.001, and #### p < 0.0001 vs HIE; $$$ p < 0.001, and $$$$ p < 0.0001 vs HIE+N-MSC.

At P21, HIE animals displayed significant impairment in object recognition, with a discrimination ratio of 39.45 ± 4.3 %, significantly lower than controls (68.38 ± 2.63 %, p < 0.0001). Animals in HIE+SSH-MSC group presented a substantial recovery (72.75 ± 6.07 %, p < 0.0001 vs. HIE), with recognition ratios comparable to control animals. Similarly to what was observed in the motor tests, animals from the HIE+N-MSC and HIE+MH-MSC groups did present improvement (42.65 ± 3.61 % and 48.47 ± 3.69 %, respectively), with a performance significantly below the HIE+SSH-MSC group (p < 0.0001 and p = 0.0011, respectively). Recognition memory impairments in all the groups subjected to HIE lesion were more notorious at P38. Moreover, the beneficial effects of administering SSH-preconditioned UC-MSCs were maintained 26 days after administration (i.e., P38).

Across all behavioral parameters, SSH-UC-MSC demonstrated superior therapeutic efficacy compared to both MH-and N-UC-MSC, highlighting the potential of SSH preconditioning in enhancing the therapeutic efficacy of UC-MSCs.

### 2.2. Proteomic insights into the recovery mechanisms driven by administration of hypoxia preconditioned-UC-MSCs

To gain insight into the recovery mechanisms occurring in animals treated with IV administration of UC-MSCs, proteomic analysis was performed using SWATH/DIA-MS on ipsilesional hemisphere brain tissue collected from control, lesioned animals (HIE), and lesioned animals treated with IV-administration of naïve UC-MSCs (HIE+N-MSCs) or UC-MSCs exposed to 0.1% oxygen for 24 hours (HIE+H-MSCs) at P40.

#### 2.2.1. Impact of UC-MSC Treatment on Protein Expression After Neonatal HI Injury

A library of 2,693 proteins was constructed with DDA mode using the rat’s brain tissue collected at P40 (30 days post-injury and 28 days post-treatment, divided into ipsilesional and contralesional hemispheres). SWATH/DIA-MS analysis enabled the quantification of 1,923 proteins across all experimental conditions. After determining the peak area of each fragment and its precursor ion with PeakView (Sciex), the unique peptides were quantified by the sum of total peak area of the precursor ion. The total peak area of each protein was calculated by the sum of its peptides. Then, proteins’ quantification was normalized to the total sum of all proteins present in the experimental sample. Peptide relative peak areas were normalized for the total sum of fragment areas for the respective sample. Further analysis identified 562 proteins differentially expressed among the experimental groups (p < 0.05 with Kruskal-Wallis test AND VIP score > 1), indicating alterations resulting from HI injury and/or subsequent UC-MSCs’ administration (Figure 3A).

**Figure 3.**
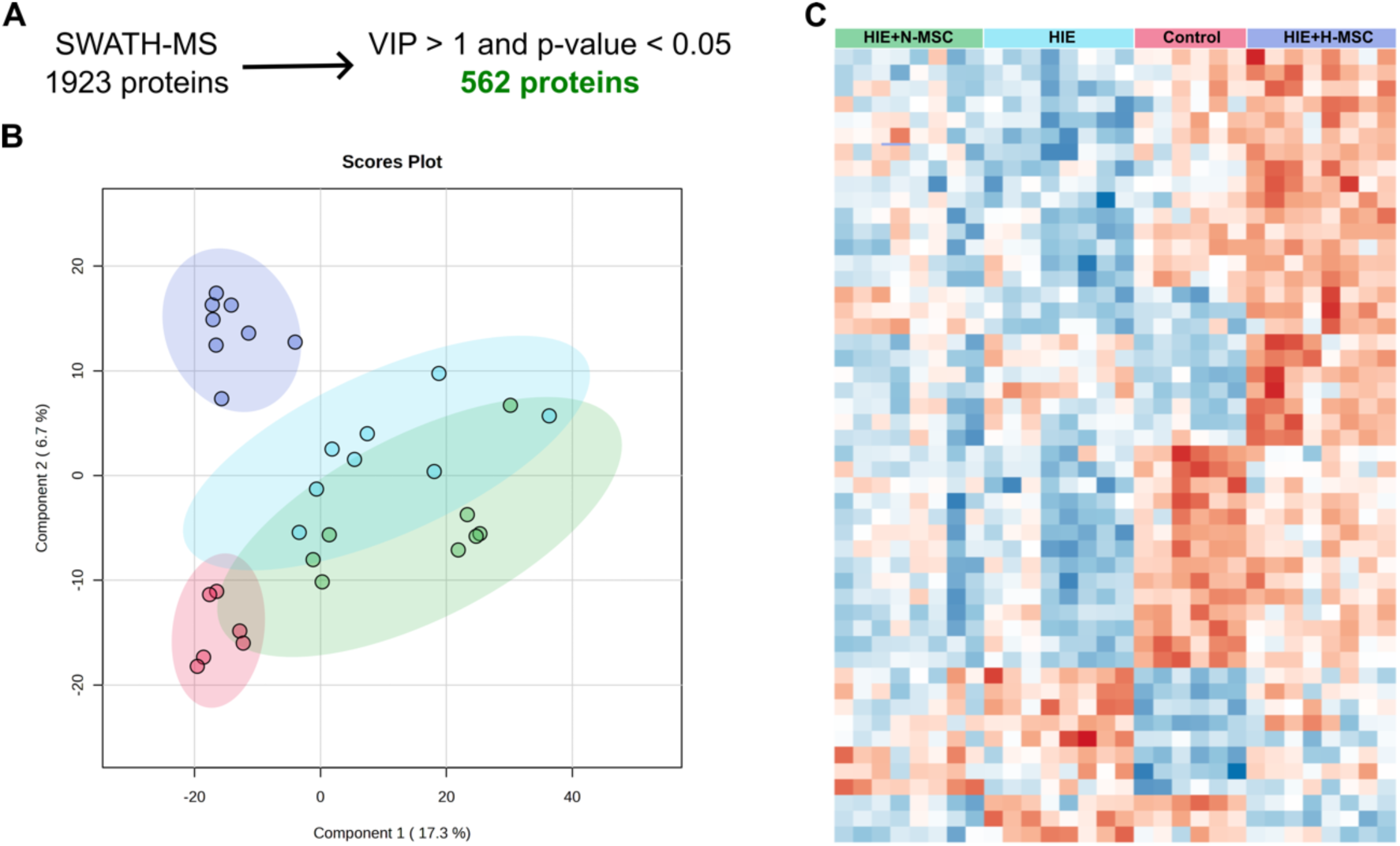
Proteomic profile of rats’ brain tissue following neonatal HI injury and administration of naïve UC-MSCs or hypoxia-preconditioned UC-MSCs. (A) Selection of SWATH/DIA-MS proteins to identify differentially expressed proteins based on statistical significance (Kruskal-Wallis test) and VIP score greater than one. (B) Partial least squares-discriminant analysis (PLS-DA) of the 1923 proteins quantified across all experimental groups. (C) Heatmap depicting expression changes of the top 50 VIP proteins identified by PLS-DA across the four experimental groups. Legend: Red – Control; Light blue -HIE; Green – HIE+N-MSC; Dark blue – HIE+H-MSC.

PLS-DA of the quantified proteome revealed distinct clustering patterns among the experimental groups (Figure 3B). HI injury induced a clear shift in the molecular profile, with HIE and HIE+N-MSC groups clustering together. The clustering of HIE and HIE+N-MSC groups indicates that naïve UC-MSC treatment did not substantially alter the HI-induced molecular changes. Hypoxia-preconditioned UC-MSC (H-MSC) IV administration in HIE animals generated a unique molecular signature, distinctly separated from both the HIE/HIE+N-MSC cluster along both components 1 and 2. The HIE+H-MSC group was closer to control, although it was separated by component 2 of the analysis. These observations suggest that animals that had IV administration of H-MSC had a partial recovery of brain function, with some pathways being uniquely triggered by the administration of these cells after injury.

Hierarchical clustering analysis of the 50 proteins with the highest VIP scores supported the distinct expression patterns that characterized each experimental condition (Figure 3C). First, samples were correctly attributed to the respective experimental group without supervised clustering, indicating consistent and reproducible effects caused by the lesion and UC-MSCs’ administration. Moreover, the HIE+H-MSC group demonstrated a protein expression profile resembling control conditions more closely than other treatment groups.

#### 2.2.2. Overrepresented gene ontologies in the ipsilesional hemisphere 30 days after neonatal HI injury

A comprehensive pathway analysis of the 562 DEPs identified through proteomic screening analysis revealed an extensive modulation of crucial neural pathways and cellular components among the different experimental groups.

Pathway enrichment analysis using Reactome, an open-source online database of biological pathways, highlighted significant alterations in neural signaling and protein synthesis pathways (Figure 4A and B). Pathways related to synaptic transmission and overall neuronal system function were substantially enriched. Gene ontology analysis of biological processes revealed enrichment in pathways associated with energy metabolism and synaptic function (Figure 4C). Specifically, pathways involving fatty acid beta-oxidation and cellular respiration showed significant representation, indicating a potential restoration of energy metabolism.

**Figure 4.**
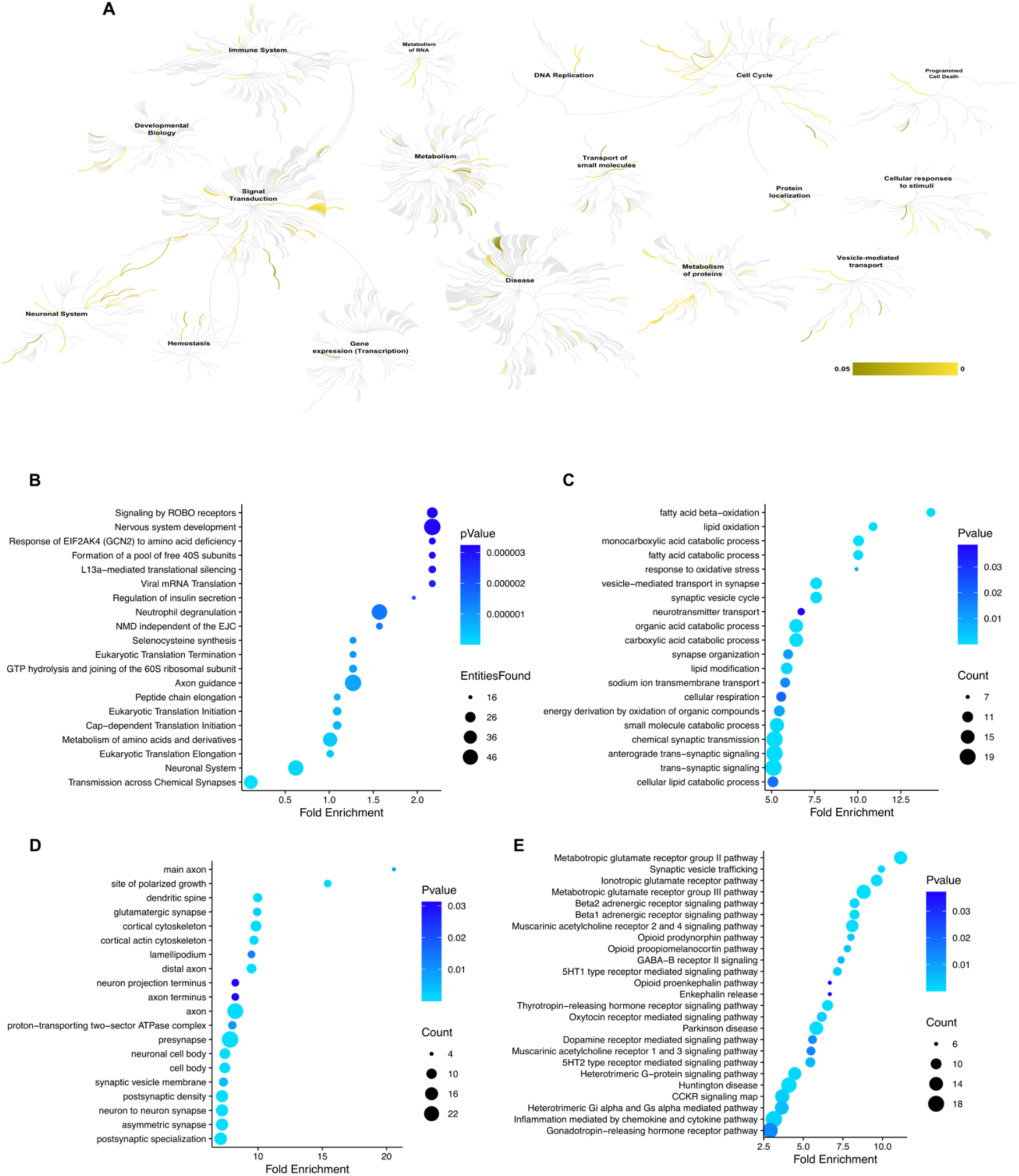
Pathways and gene ontologies in the differentially expressed proteins in ipsilesional hemisphere of lesioned animals or animals treated with UC-MSCs. (A, B) Overall enriched pathways and top 20 overrepresented enriched Reactome pathways. (C-E) Enriched gene ontologies for biological processes, cellular component, and PANTHER pathways.

Gene ontology analysis of cellular components further revealed extensive remodeling of neuronal structures (Figure 4D). Proteins linked to synaptic structures were particularly enriched, including presynaptic components and postsynaptic density proteins. Axonal components, including main axons and distal axons, were also significantly enriched, along with cytoskeletal proteins, suggesting ongoing neuronal reorganization. Additionally, PANTHER pathway classification analysis, an online tool that uses controlled vocabulary to describe pathways, their components, and the relationships among them, highlighted significant enrichment in neurotransmitter signaling pathways (Figure 4E). Glutamatergic signaling pathways, including metabotropic and ionotropic glutamate receptors, had marked enrichment. Modulatory neurotransmitter systems, including adrenergic, GABAergic, and serotonergic pathways, were also enriched.

#### 2.2.3. Long-term effects of hypoxia-preconditioned UC-MSCs administration on protein expression

To determine the impact of injury and/or UC-MSC administration on the up-or-down-regulation of the 562 DEPs, we assessed both shared and unique DEPs across the experimental groups (Figure 5A).

**Figure 5.**
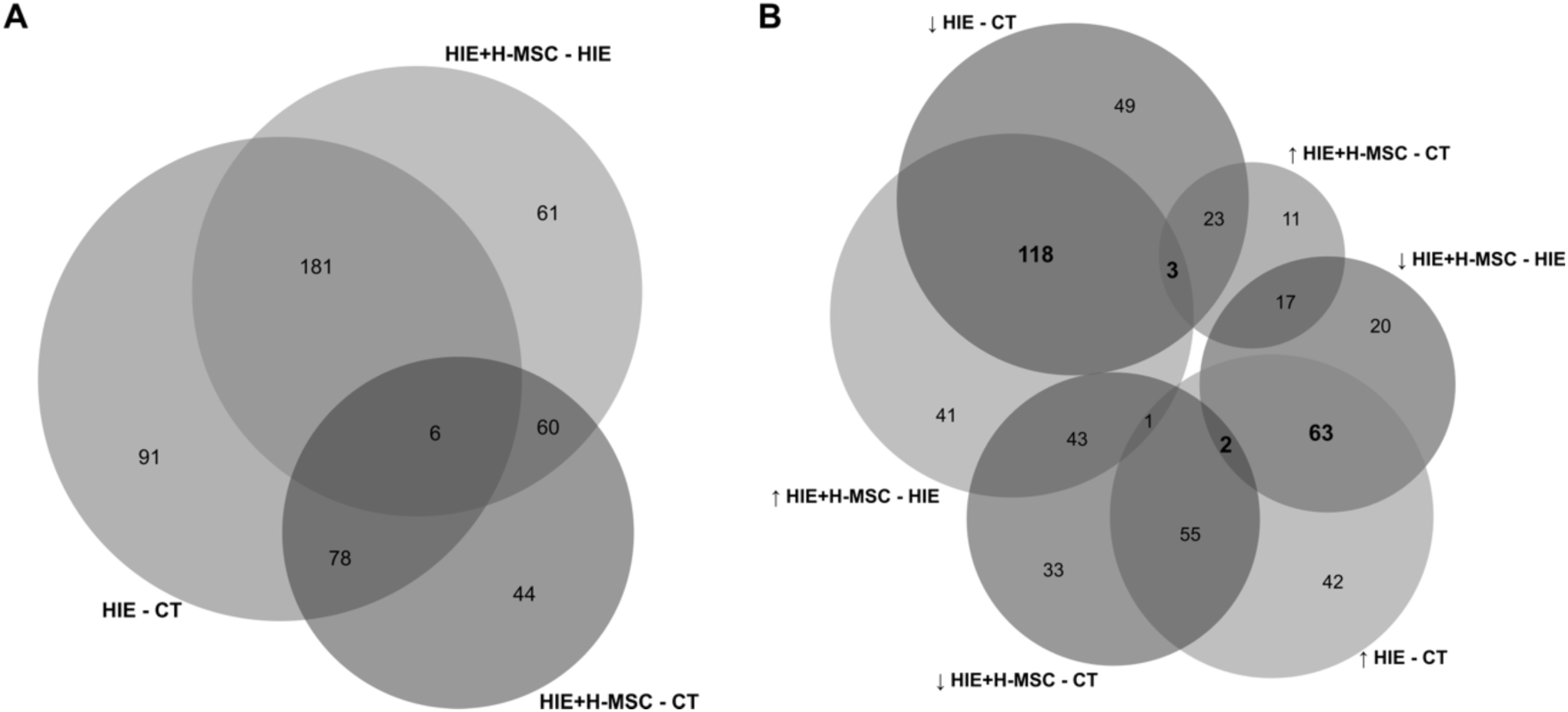
Differential protein expression, across experimental groups, 30 days after neonatal HI brain injury. (A) Venn diagram with DEPs across experimental groups. (B) Venn diagram illustrating the intersection between proteins regulated by HIE and/or H-MSC administration.

In the HIE vs. control comparison, 91 proteins were uniquely altered, while in the HIE+H-MSC vs. control comparison 44 proteins were uniquely altered. In the HIE+H-MSC vs. HIE comparison, 61 proteins exhibited unique regulation, indicating a long-term effect of the molecular alterations that were triggered by H-MSCs administration. Among the 187 DEPs shared between the HIE and HIE+H-MSC groups, 65 proteins were upregulated in the HIE group but downregulated in the HIE+H-MSC group and 121 proteins downregulated in HIE were upregulated in the HIE+H-MSC group (Figure 5B), indicating that hypoxia-preconditioned UC-MSC administration might reverse certain injury-induced changes.

#### 2.2.4. Protein-protein interaction networks 30 days post-injury and after treatment with hypoxia-preconditioned UC-MSCs

The PPI network analysis revealed distinct clusters of DEPs 30 days after brain injury, highlighting the complex molecular changes induced by HI damage that remain in later phases of injury and/or that were modulated by the administration of hypoxia-preconditioned UC-MSCs. Using the STRING bioinformatics platform set at a high confidence level and Markov Cluster Algorithm clustering (inflation parameter 1.5), multiple functional groups were identified within the DEPs. The largest group of downregulated proteins was associated with the postsynaptic density (32 proteins) (Figure 6), suggesting impaired synaptic structure and function in the HIE brain.

**Figure 6.**
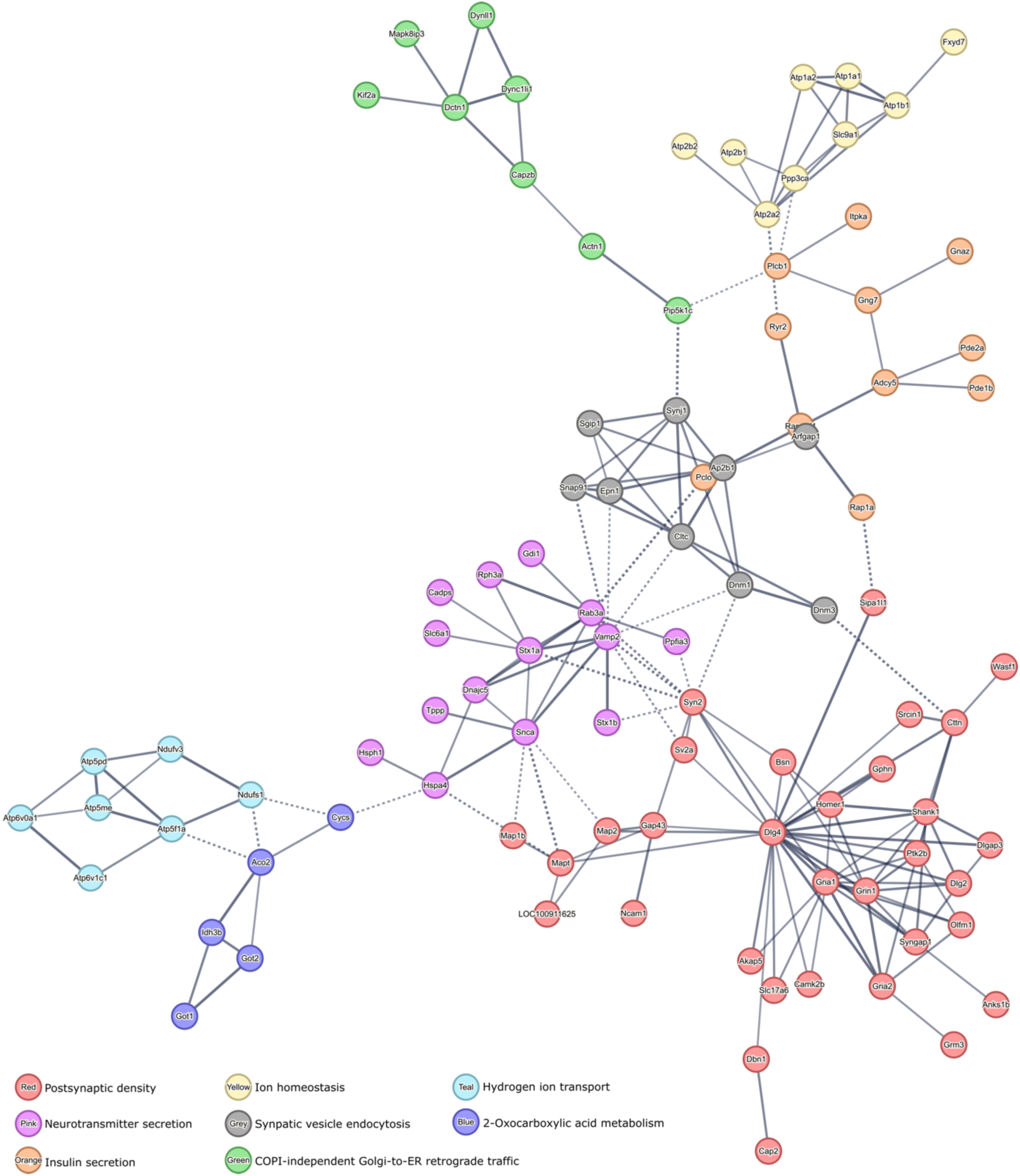
Protein-protein interactions with more than five downregulated proteins in the ipsilesional hemisphere of HIE animals. High confidence PPI was identified by MCL clustering (1.5 inflation parameter) with the STRING bioinformatics tool. Network nodes represent proteins and edges represent protein-protein associations. Thickness of protein-protein associations reflects the level of confidence.

This was followed by pathways involved in neurotransmitter secretion (14 proteins) and insulin secretion (11 proteins), both crucial for maintaining neuronal communication and metabolic regulation, respectively. The downregulation of proteins involved in these pathways highlights potential disruptions in synaptic signaling and energy homeostasis following hypoxic injury. Additionally, proteins linked to ion homeostasis (9 proteins) and synaptic vesicle endocytosis (9 proteins) were also downregulated, further underscoring potential deficiencies in maintaining neuronal excitability and neurotransmitter recycling. Several other pathways were identified with fewer proteins but still indicative of altered neuronal and metabolic processes, such as COPI-independent Golgi-to-ER retrograde traffic (8 proteins), hydrogen ion transport (7 proteins), and 2-oxocarboxylic acid metabolism (5 proteins).

In line with the observed downregulation of key proteins involved in synaptic and metabolic pathways, the analysis of upregulated proteins in the HIE brain 30 days post-injury revealed a complementary set of molecular alterations (Figure 7). The most prominent upregulated cluster was associated with the ribosome (23 proteins), suggesting an increased demand for protein synthesis during the chronic phase of injury. This likely reflects an effort to promote cellular repair and regeneration processes. In addition, proteins involved in fatty acid degradation (18 proteins) were upregulated, pointing toward enhanced lipid metabolism as a response to cellular stress and energy needs. Detoxification pathways (9 proteins) were also upregulated, highlighting the brain’s attempt to counteract oxidative stress and damage caused by the hypoxic environment. Other pathways include axonal growth stimulation (6 proteins), protein import (6 proteins), and calcium ion binding involved in the regulation of presynaptic cytosolic calcium ion concentration and astrocyte end-feet (6 proteins), indicating an active effort to promote neuronal repair and regulate intracellular calcium levels, which are critical for synaptic function and signaling.

**Figure 7.**
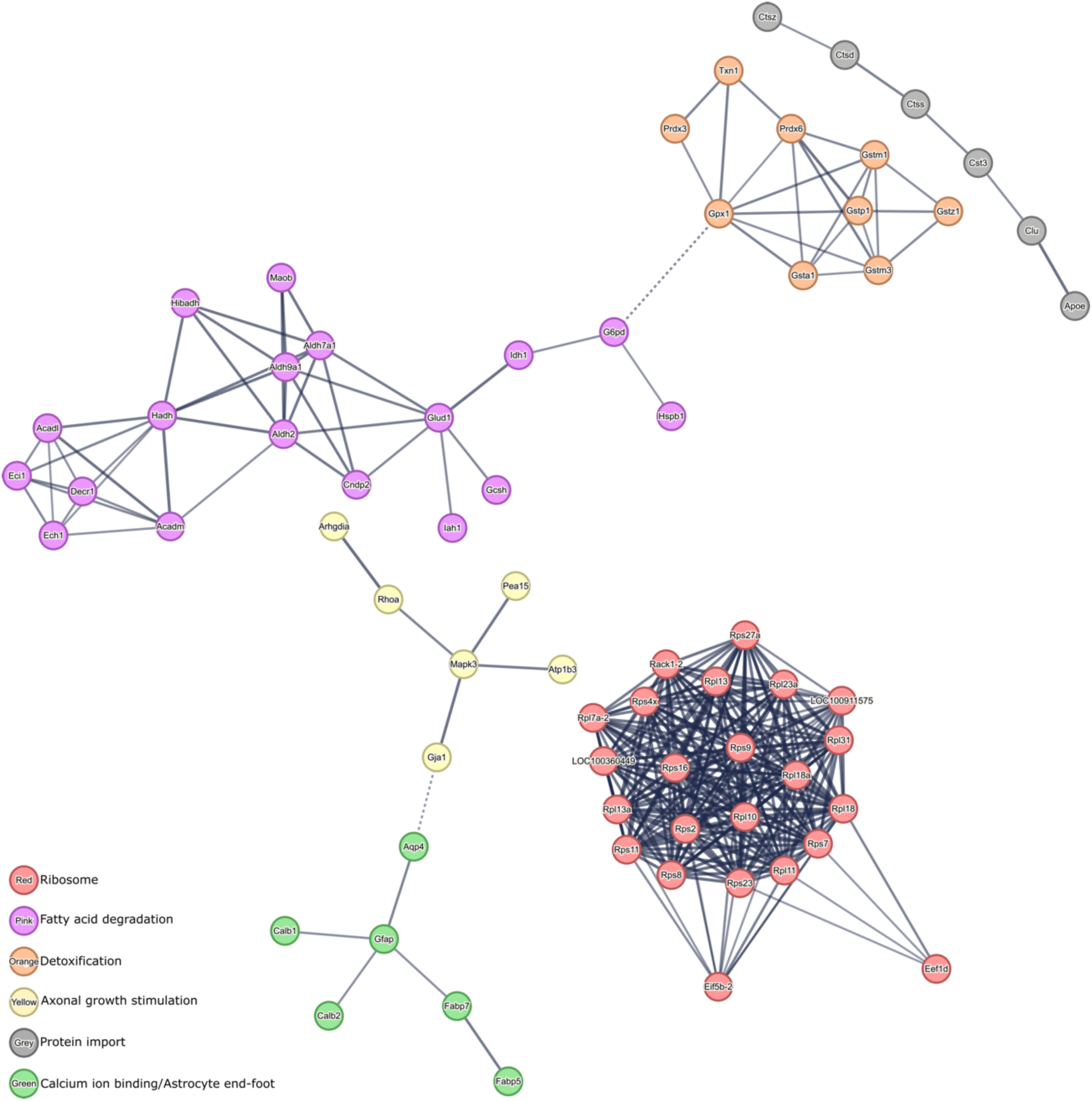
Protein-protein interactions with more than five upregulated proteins in the ipsilesional hemisphere of HIE animals. High confidence PPI was identified by MCL clustering (1.5 inflation parameter) with the STRING bioinformatics tool. Network nodes represent proteins and edges represent protein-protein associations. Thickness of protein-protein associations reflects the level of confidence.

The analysis of PPI networks formed by proteins that were differentially regulated between HIE animals and H-MSCs-treated animals allowed to identify pathways that most likely were modulated by the effects of H-MSCs administration in lesioned animals (Figure 8).

**Figure 8.**
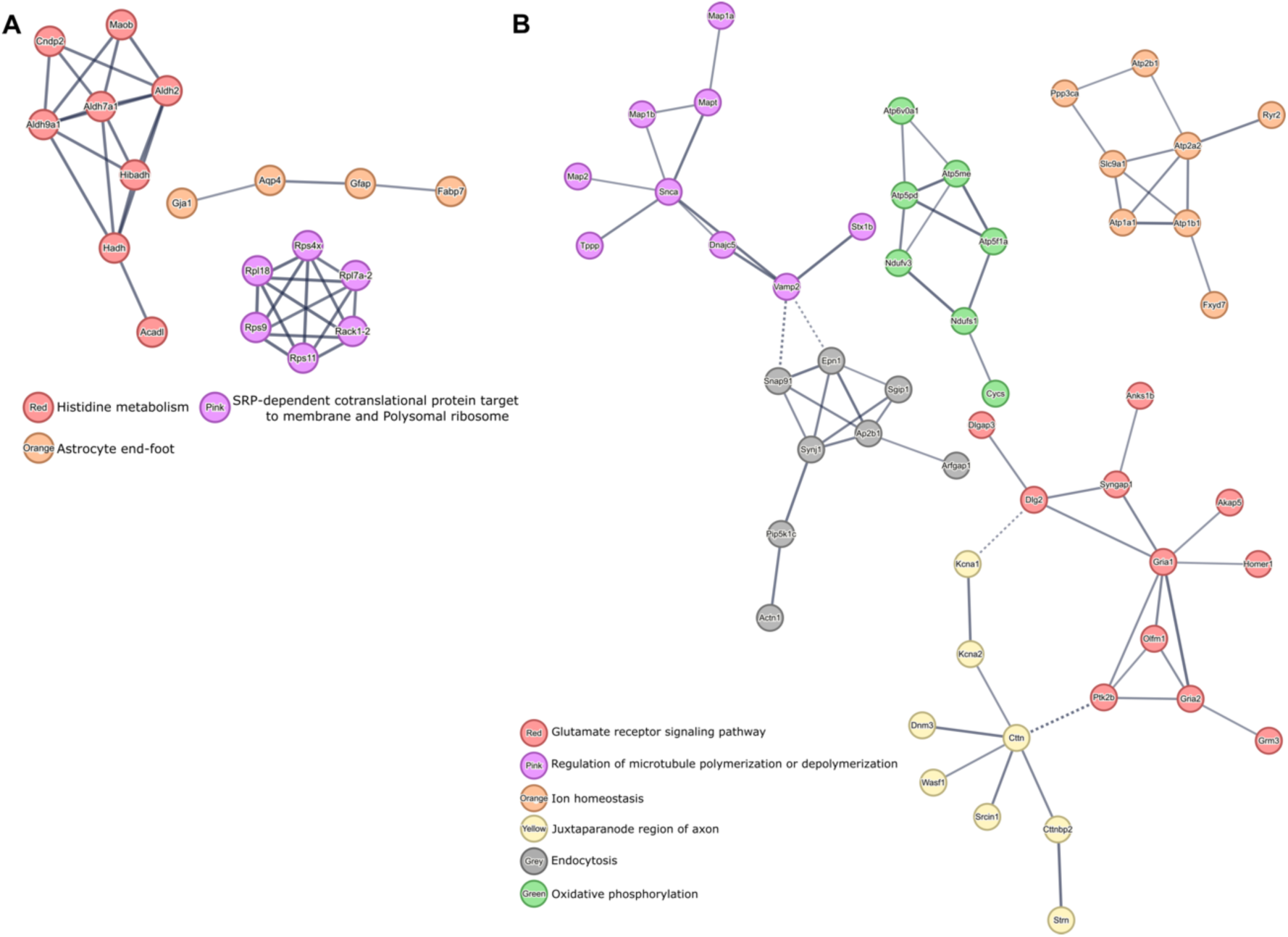
Protein-protein interactions formed by DEPs between HIE and HIE+H-MSC animals. (A) PPI of upregulated proteins in HIE animals that were downregulated in HIE+H-MSC animals and (B) PPI of downregulated in HIE animals that were upregulated in HIE+H-MSC animals. High confidence PPI was identified by MCL clustering (1.5 inflation parameter) with the STRING bioinformatics tool. Network nodes represent proteins and edges represent protein-protein associations. Thickness of protein-protein associations reflects the level of confidence.

The PPI networks formed by proteins upregulated in HIE animals but downregulated in HIE+H-MSC animals (Figure 8A) were related to histidine metabolism (8 proteins) and SRP-dependent cotranslational protein targeting to the membrane (6 proteins), and astrocyte end-foot function (4 proteins), underscoring the astroglia-remodeling occurring during injury that is no longer overrepresented in H-MSC-treated animals.

On the other hand, analysis of PPI networks formed by downregulated proteins in HIE that were upregulated in HIE+H-MSC animals (Figure 8B) revealed a significant involvement of the glutamate receptor signaling pathway (11 proteins). Furthermore, pathways related to microtubule regulation (9 proteins) and ion homeostasis (8 proteins) indicate disruptions in cellular stability and homeostasis that were recovered in animals treated with H-MSC. Also, the identification of proteins of the juxtaparanode region of the axon (8 proteins) and endocytosis pathway (8 proteins) suggest challenges in maintaining axonal integrity and neurotransmitter recycling post-injury. Also, PPI analysis revealed a potential long-term upregulation of proteins involved in oxidative phosphorylation (7 proteins) in animals of the HIE+H-MSCs group.

In the search for pathways that become modulated by the changes induced by the H-MSCs administration in lesioned animals, proteins that were differentially expressed only between control and HIE+H-MSCs groups were also analyzed (Figure 9). The PPI networks formed by these proteins were related to circadian entrainment (8 proteins), fibrinolysis and acute phase response (6 proteins), and the respiratory chain (5 proteins). A common feature among these networks is their involvement in essential regulatory processes that support recovery and homeostasis, including stress response, energy production, and the modulation of inflammation and repair mechanisms.

**Figure 9.**
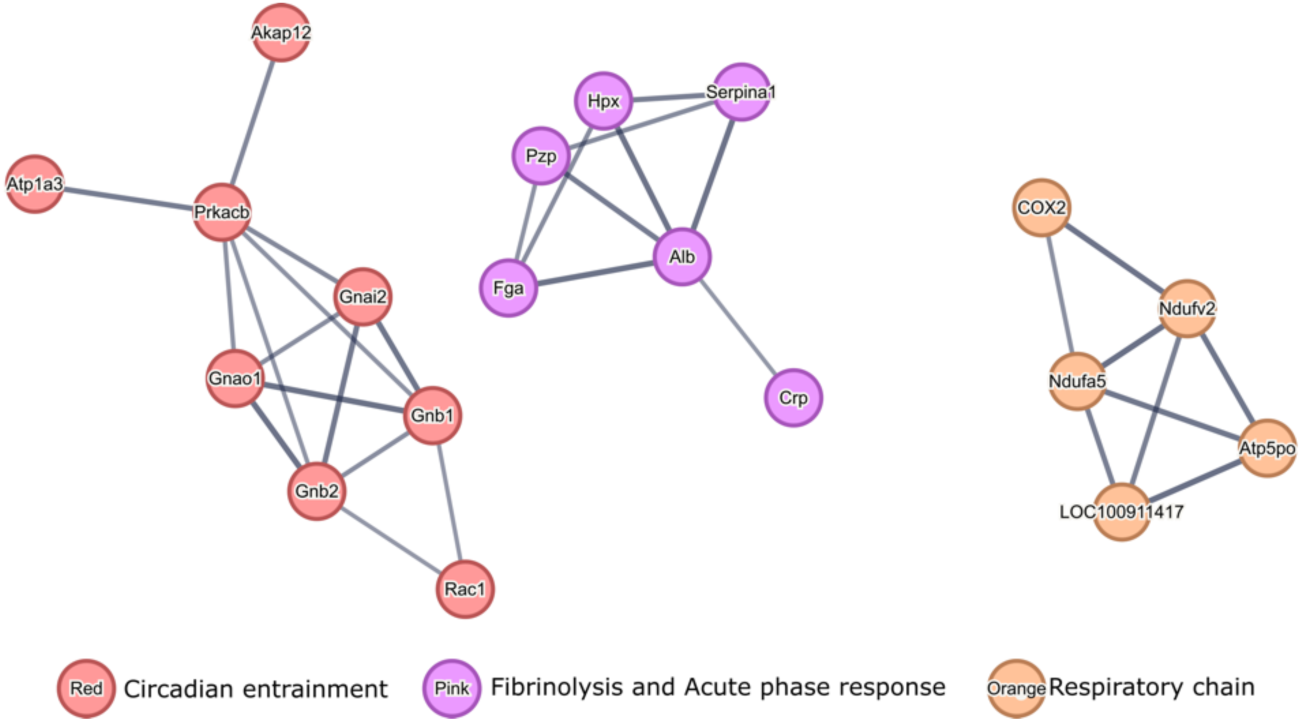
Protein-protein interactions formed by proteins differentially expressed between control animals and HIE+H-MSC animals. High confidence PPI was identified by MCL clustering (1.5 inflation parameter) with the STRING bioinformatics tool. Network nodes represent proteins and edges represent protein-protein associations. Thickness of protein-protein associations reflects the level of confidence.

#### 2.2.5. Characterization of protein expression regulation 30 days after brain injury in animals treated with hypoxia-preconditioned UC-MSCs

To further explore the differences between the proteome of HIE animals and those treated with UC-MSCs, naïve or preconditioned with hypoxia. The proteins showing differential expression between Control and HIE (HIE – CT), HIE+H-MSC and Control (H-MSC – CT), HIE+H-MSC and HIE (H-MSC – HIE), HIE+N-MSC and Control (N-MSC – CT), and HIE+N-MSC and HIE (N-MSC – HIE) were analyzed with the PANTHER tool to identify the top 15 overrepresented biological processes in each of these comparisons. To facilitate the visualization of these processes we divided them into six categories: synaptic function and neural communication (Figure 10A), intracellular transport and structural organization (Figure 10B), metabolism and oxidative processes (Figure 11A), cellular detoxification and redox homeostasis (Figure 11B), ion transport and homeostasis (Figure 11C), and neurodevelopment and growth (Figure 11D).

**Figure 10.**
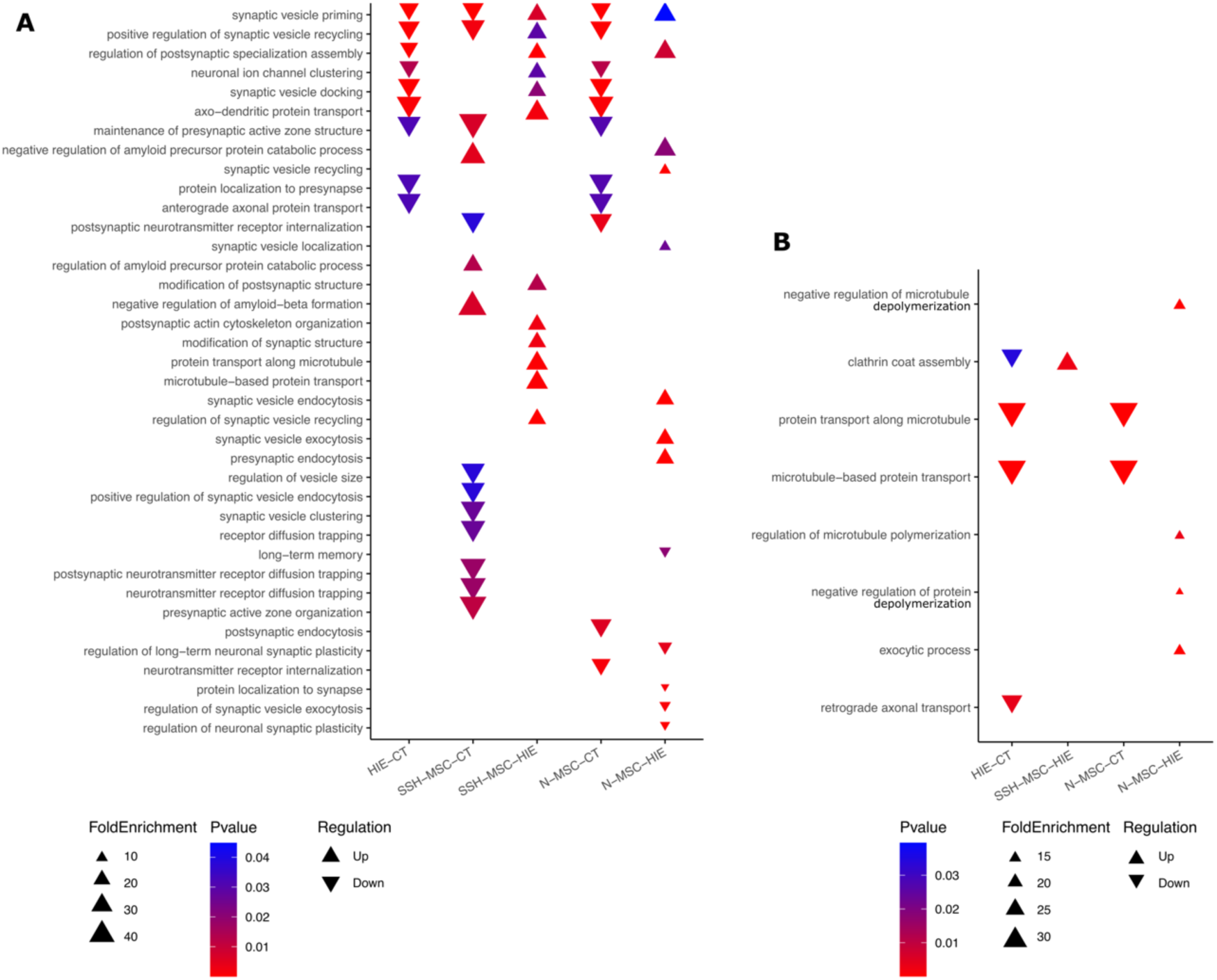
Overrepresented gene ontologies for top 15 biological processes involved in (A) synaptic function and neural communication and (B) intracellular transport and structural organization in differentially expressed proteins amongst the experimental groups control, HIE, HIE+H-MSC and HIE+N-MSC.

**Figure 11.**
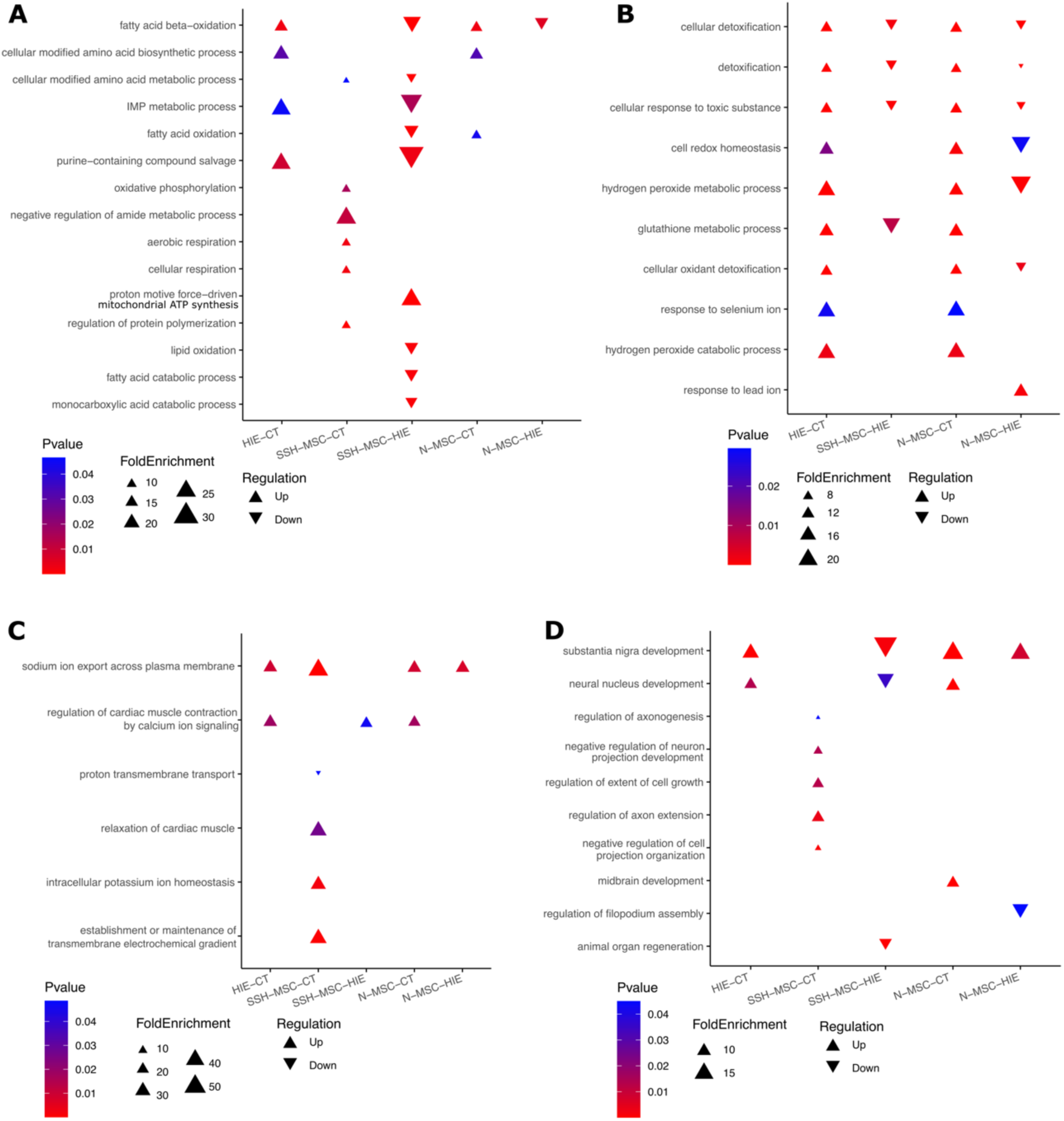
Overrepresented gene ontologies for top 15 biological processes involved in (A) metabolism and oxidative processes, (B) cellular detoxification and redox homeostasis, (C) ion transport and homeostasis, and (D) neurodevelopment and growth in differentially expressed proteins amongst the experimental groups control, HIE, HIE+H-MSC and HIE+N-MSC.

Gene ontology revealed that HIE animals had downregulation of synaptic-related processes (Figure 10A), when compared with the control group, suggesting impaired neuronal communication and synaptic integrity even 30 days after the lesion induction. Lesioned animals treated with H-MSCs, when compared to control, still had some of these processes downregulated, however, the biological processes related to synaptic vesicle priming, synaptic vesicle recycling, synaptic vesicle docking, postsynaptic assembly, neuronal ion channel clustering, and axo-dendritic protein transport where upregulated when compared with HIE non-treated animals (Figure 10A, first six processes). Nonetheless, when analyzing the tissue of animals from HIE+N-MSC group, it was found downregulation of the several synaptic processes compared to control and less representation of processes that were upregulated in HIE+H-MSC animals compared to HIE group (Figure 10A). Moreover, HIE and HIE+N-MSC groups showed a downregulation of biological processes related to intracellular transport and structural organization, which was not found in HIE+H-MSC animals (Figure 10B). The upregulation of these processes in HIE+H-MSC group versus HIE group suggests a potential recovery in axonal transport and synaptic function, which contribute to the restoration of neuronal connectivity and communication.

In contrast, the DEPs between HIE and HIE+H-MSC groups revealed a downregulation of biological pathways related with metabolism and oxidative processes as well as cellular detoxification and redox homeostasis (Figure 11A, B). The gene ontology of DEPs in the HIE+N-MSC group revealed overlap with the HIE+H-MSC group in reduced detoxification pathways (Figure 11C). Additionally, all groups had upregulation in neurodevelopmental processes, such as substantia nigra development and neural nucleus development, when compared with control (Figure 11D). Only HIE+H-MSC group had a downregulation of these processes when compared with untreated lesioned-animals.

These results also revealed that animals treated with naïve UC-MSCs lacked significant enrichment in axonal and synaptic transport processes upregulated in HIE+H-MSC animals, suggesting a therapeutic advantage of hypoxia-preconditioned UC-MSCs in promoting recovery through enhanced protein transport and neuronal connectivity.

### 2.3. HIE induces long-term glial changes that are reduced by hypoxia-preconditioned UC-MSCs

Given the central role of glial cells in both HIE injury progression and recovery, we examined whether hypoxia-preconditioned UC-MSCs modulated long-term glial responses in the HIE rat model. By exploring this dimension, we aimed to understand if hypoxia-driven preconditioning not only enhances functional recovery but also impacts the cellular environment of the injured brain, potentially contributing to sustained neuroprotection and repair.

The levels of proteins associated with glial cells function and reactivity was analyzed to better understand the contribution of these cells to the long-term effects of HI injury as well as the impact of hypoxia preconditioned-UC-MSCs’ IV administration. Our previous work demonstrated that HIE induces significant glial reactivity, with a marked increase in GFAP and Iba1 expression and alterations in astrocyte and microglia morphology (Chapter 4). Building on this foundation, this study explored other glial-associated proteins to assess their differential expression profiles 30 days post-HI injury, as measured by SWATH/DIA-MS. The proteins that were selected for analysis included aquaporin-4 (AQP4), apolipoprotein E (APOE), CD44 antigen, chloride intracellular channel protein 1 (CLIC1), fatty acid-binding protein, brain (FABP7), GFAP, and syndecan-4 (SDC4). These proteins are known to be involved in various glial cell functions such as water homeostasis (AQP4), cholesterol transport (APOE), extracellular matrix interactions (CD44), and ion transport (CLIC1), all of which are crucial for glial-mediated responses (Figure 12).

**Figure 12.**
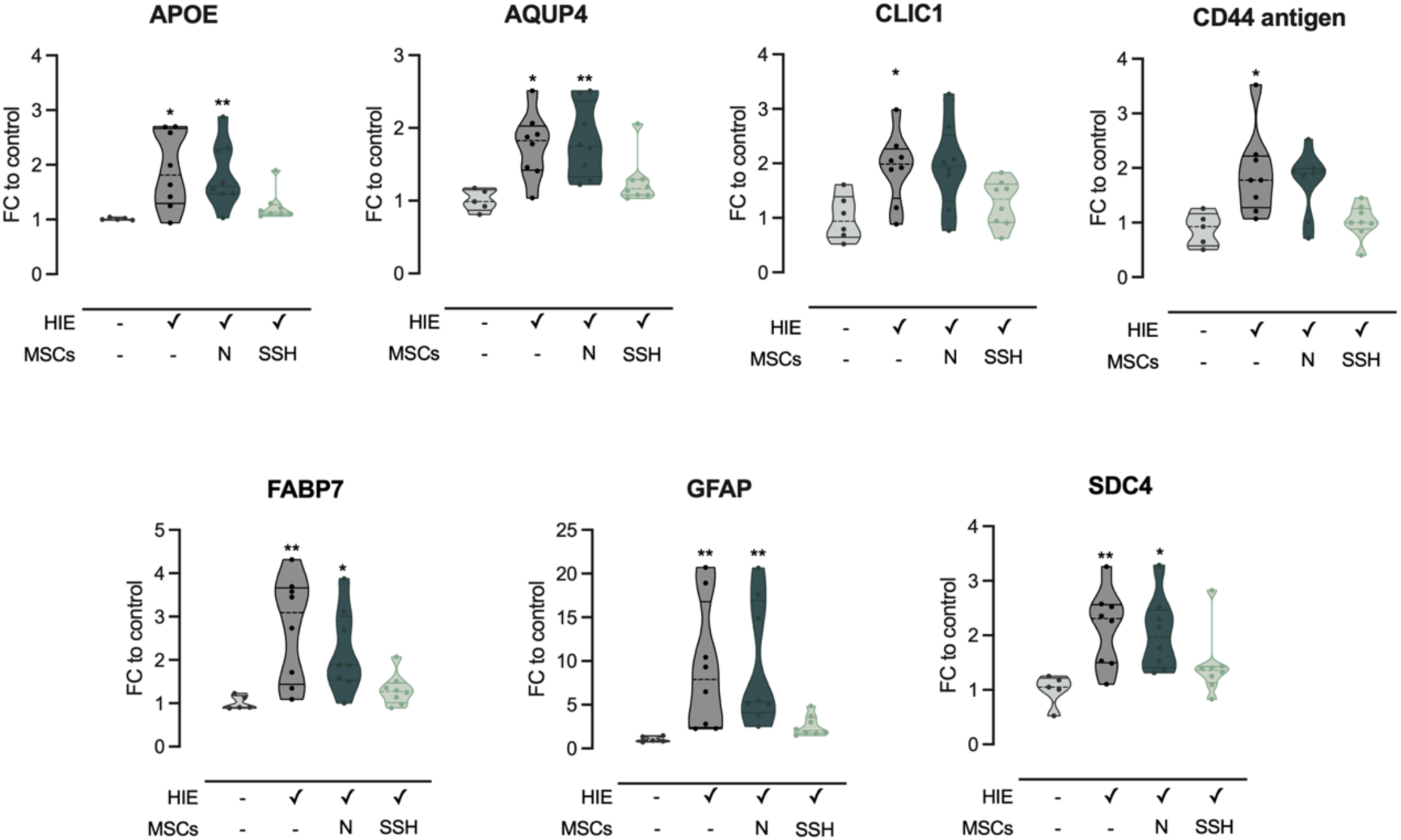
Changes in glial cell-related proteins 30 days after HIE injury. Proteins were quantified by SWATH/DIA-MS and statistical analysis was performed using the Kruskal–Wallis test and Dunn’s multiple comparison test. Statistical differences between groups are represented in the figures with * p < 0.05 and ** p < 0.01 compared to control. Data expressed as fold-change (FC) to control. Legend: AQP4 – aquaporin 4; APOE – apolipoprotein E; CD44 – CD44 antigen; CLIC1 – chloride intracellular channel protein 1; FABP7 – fatty acid binding protein, brain; GFAP – glial fibrillary associated protein; SDC4 – syndecan-4.

As expected, GFAP, a hallmark of astrocyte reactivity (37), was substantially upregulated in HIE animals compared to controls. A significant 9.15-fold increase (p = 0.0023) was observed in GFAP levels in animals of HIE group, corroborating the extensive reactivity previously described in the literature. GFAP expression remained elevated only in the N-MSCs-treated group (9.4-fold increase, p = 0.0011), with HIE+SSH-MSCs animals no longer presenting upregulation of this protein. AQP4, the main water channel in the CNS, and present in astrocyte end-feet (38), also exhibited 1.76 fold increase of its expression post-HI injury (p = 0.0124). HIE+N-MSC animals, but not animals treated with SSH-preconditioned UC-MSCs, displayed an increase in AQP4 levels (1.82-fold, p = 0.0087). APOE, a protein that regulates lipid transport from neurons to glial cells under HI conditions (39), was also upregulated in HIE animals (1.90-fold, p = 0.0106). In the treatment groups, only HIE+SSH-MSCs animals presented APOE levels closer to control (1.24-fold increase compared to control), though the differences with the HIE group were not statistically significant. Both CD44, a cell surface glycoprotein that acts as a receptor for hyaluronic acid, and SDC4, a transmembrane heparan sulfate proteoglycan involved in cell adhesion, exhibited increased expression following HI injury (1.90-fold, p = 0.0386 and 2.14-fold, p = 0.0087, respectively). Treatment with SSH-preconditioned UC-MSCs normalized the levels of these proteins to control levels.

CLIC1, a protein involved in regulating chloride ion transport in astrocytes and microglia (40, 41) and potential marker of inflammatory reactive astrocytes (42), had almost a 2-fold increase in HIE animals (1.92-fold, p = 0.0365). Like for the other glial cells-related proteins, the HIE+SSH-MSC group showed a tendency toward normalized CLIC1 levels, although the changes were not statistically significant from HIE.

Finally, FABP7, a marker of neural stem cells and astrocytes (43), was elevated in HIE animals (2.74-fold, p = 0.0068). Although the administration of naïve UC-MSCs was incapable of decreasing FABP7 expression (2.19-fold to control, p = 0.0268), the administration of UC-MSCs preconditioned with SSH reduced its expression to control levels.

## 3. Discussion

The results presented here demonstrate the potential of hypoxia-preconditioned UC-MSCs as a neuronal repair strategy after HI insult to the developing brain. Using proteomic profiling, pathway enrichment analyses, as well as motor and cognitive evaluations we have shown that SSH-preconditioned UC-MSCs mitigated injury-induced impairments and promoted recovery, closely resembling the physiological state of control animals.

The behavioral assessments suggested animals treated with UC-MSCs preconditioned with SSH had a superior neurological recovery compared to animals treated with MH (5% oxygen, 48 hours) or naïve MSCs. Indeed, the therapeutic advantage of administering hypoxic-preconditioned UC-MSCs has been observed in several preclinical models of conditions affecting the CNS, such as stroke, TBI and SCI (15). Most studies used MSCs derived from various sources, such as BM, adipose tissue, and UC, thus suggesting that this strategy could be applied to MSCs independently of their source.

A review by Samal et al., 2021, highlighted multiple factors, including oxygen concentration and exposure duration, that can influence MSC phenotype under hypoxic conditions (31). Our findings align with this finding, as short-term, severe hypoxia (0.1% oxygen for 24 hours) was more effective in reducing motor and cognitive impairments in HIE animals than MH. This suggests that the therapeutic potential of MSCs is strongly influenced by the specific hypoxic preconditioning protocol used. MH supports sustained growth, but short, severe hypoxia seems to trigger a more robust adaptive response, possibly leading to better therapeutic outcomes. This differential response is likely due to distinct molecular mechanisms: severe hypoxia can rapidly increase HIF-1α levels and its nuclear translocation, while mild hypoxia can activate HIF-2α (35, 44). Acute hypoxic exposure is likely to create a "priming effect," allowing MSCs to mount a maximal adaptive response without the detrimental effects of prolonged severe hypoxia. This is evidenced by increased Akt phosphorylation under short-term severe hypoxia, promoting the activation of cell survival pathways (36). Additionally, the activation of NF-κB by short-term hypoxic exposure leads to the upregulation of growth factors like VEGF, FGF2, HGF, and insulin-like growth factor (IGF) (45). The hypoproliferative state observed under severe hypoxia suggests a conservation of energy, likely mediated by the mechanistic target of rapamycin complexes 1 and 2, to optimize energy metabolism and stress resistance (46). These molecular adaptations may explain why the administration of MSCs preconditioned with SSH resulted in more pronounced functional improvements compared to the administration of MSCs exposed to prolonged MH in the HIE rodent model.

On the other hand, the molecular response of MSCs to hypoxia appears to be centrally regulated by prolyl hydroxylase domain proteins. These proteins hydroxylate key signaling molecules like HIFs, NF-κB, and protein kinase B (Akt), in an oxygen-dependent manner (47). Under normoxic conditions, prolyl hydroxylase domain proteins hydroxylate HIF-α, leading to its proteasomal degradation, while in hypoxia, the lack of hydroxylation allows HIF-α to accumulate, translocate to the nucleus, and activate hypoxia-related genes (48). Similarly, hypoxia prevents the hydroxylation of NF-κB and Akt, thus blocking their inactivation and promoting cell survival (49). Previous studies have looked for the impact of hypoxia on the proteome and secretome of MSCs to understand better what drives this enhanced therapeutic effect. For instance, hypoxia preconditioning enhanced the survival and migration ability of MSCs both *in vitro* and after *in vivo* administration (50–52). Moreover, hypoxia-preconditioned MSCs also have increased homing capacity. This stimulus upregulated the expression of stromal cell-derived factor 1 receptors, such as CXC chemokine receptor 4, 7 and CX3C motif chemokine receptor 1 (28, 32), which are crucial for the migration of stem cells towards injury sites (53). Indeed, *in vivo* studies have demonstrated that hypoxic-preconditioned MSCs had better survival, migration and homing abilities than normoxic MSCs in rodent models of cerebral ischemia (13, 17, 21, 54). These observations were also accompanied by improved neurological function.

Proteomic profiling of ipsilesional brain tissue highlighted distinct mechanisms associated with UC-MSC treatment. Notably, the ipsilateral hemisphere of animals treated with SSH-MSCs had a proteomic profile closer to that of controls and markedly different from the HIE animals and animals treated with naïve UC-MSCs. Pathway enrichment analyses underscored key processes, such as synaptic remodeling, neurotransmitter system recovery, and metabolic reprogramming.

Upregulated pathways in HIE animals treated with the hypoxia preconditioned cells included adrenergic, GABAergic, and serotonergic pathways, suggesting an overarching restoration of neurotransmitter system function. The interaction between GABAergic, glutamatergic, and serotonergic systems plays a central role in synaptic plasticity and brain repair post-injury. Serotonergic pathways contribute to neuroplasticity, acting as a growth factor during brain development and interacting with other neurotransmitter systems like GABAergic and glutamatergic systems (55). Moreover, previous studies have shown that glutamatergic activation promotes neurogenesis and recovery post-stroke (56).

The analysis of PPI networks formed by proteins upregulated or downregulated in the HIE brain 30 days post-injury revealed a complex, multifaceted response to HI damage. Downregulated proteins were primarily associated with networks related to synaptic function and cellular signaling, including postsynaptic density, neurotransmitter secretion, and ion homeostasis. These alterations suggest a disruption in neural communication and metabolic balance following injury.

On the other hand, the upregulation of energy metabolism pathways, such as fatty acid oxidation and glycolysis, found in the brain of HIE animals was consistent with adaptive responses previously reported in perinatal asphyxia and adult stroke models, where metabolic shifts post-stroke included increased fatty acid metabolism and altered glycolysis within damaged white matter regions for prolonged periods after injury (57). Likewise, rats exposed to global perinatal asphyxia showed persistent alterations in brain energy metabolism even weeks post-insult (58).

In neonatal HIE, several molecular and metabolomic changes highlight the interplay of oxidative stress, inflammation, excitotoxicity, and mitochondrial dysfunction in its pathophysiology. We observed the upregulation of fatty acid oxidation and lipid oxidation pathways in HIE animals, which most likely contributes to mitochondrial dysfunction and oxidative stress. The neonatal brain, characterized by high concentrations of polyunsaturated fatty acids, elevated oxygen consumption, low antioxidant levels, and abundant iron, is particularly vulnerable to ROS, which modify macromolecules like lipids, proteins, and DNA, triggering inflammation and neuronal injury (59, 60). In neonatal HIE, lipid peroxidation biomarkers, such as malondialdehyde, increase over time indicating ongoing oxidative injury (61, 62). Here we report that HIE animals treated with SSH-MSCs exhibit a downregulation of fatty acid oxidation pathways and reduced lipid oxidation, suggesting diminished oxidative damage. The inefficiency of fatty acid oxidation in energy generation under hypoxic conditions, coupled with the neonatal brain’s reliance on glucose metabolism, underscores the importance of interventions that restore metabolic balance and mitigate oxidative stress. These findings, supported by observations in ischemic stroke human and animal studies (63–65), reinforce the role of lipid peroxidation and inflammation in ischemic brain injury and suggest that therapeutic modulation of these pathways could be beneficial. Nonetheless, fatty acid oxidation may have a role in sustaining brain function. In astrocytes, the metabolism of fatty acids help to maintain mitochondrial respiratory chain supercomplexes, which are less energy-efficient but promote the production of ROS at levels that are crucial for cellular signaling and cognitive processes (66). A recent study showed that, when the fatty acid oxidation is inhibited (by deleting an enzyme essential for transporting fatty acids into mitochondria), astrocytes increased the oxidation of pyruvate, which affected mitochondrial respiratory chain structure and negatively impacted cognitive functions, revealing a dual role of fatty acid metabolism in astrocytes.

Microglia and astrocytes play dual roles in HIE, contributing to both neuroinflammation and neuroprotection through mechanisms such as cytokine production, phagocytosis, and glial scar formation (67–71). Our previous work demonstrated that HIE induces significant astrocyte reactivity, with a marked increase in GFAP expression and changes in astrocyte morphology, particularly towards a more enlarged and ramified shape (72). Building on this foundation, this study explored the impact of administering preconditioned cells on glial reactivity, a key hallmark of the tertiary phase of neonatal HIE. SWATH/DIA-MS results revealed that untreated HIE brains had increased markers of glial reactivity, namely GFAP, AQP4, APOE, and FABP7, indicating the presence of neuroinflammation, gliosis, and metabolic stress that most likely contribute to impaired neurological function. The animals treated with SSH-preconditioned UC-MSCs had reduced levels of these markers, which might indicate that the glial cells of these animals presented distinct states from those of untreated HIE animals, namely a decrease in the number of glial cells with a neurotoxic profile.

## 4. Conclusion

This study demonstrates that SSH preconditioning enhanced the neurorestorative potential of UC-MSCs in neonatal HIE. Even a single, sub-optimal dose of these cells achieved significant improvements in neurological function and glial modulation, surpassing the effects of naïve or mild-preconditioned cells. Proteomic analysis revealed over 550 DEPs in the lesioned hemisphere, with intravenous administration of hypoxia-preconditioned UC-MSCs effectively reversing pathological protein expression patterns, particularly in pathways critical for synaptic function, neural development, and energy metabolism.

While this study provides valuable insights into the therapeutic potential of hypoxia-preconditioned UC-MSCs for neonatal HIE, several questions, which were not in the scope of this study, remain unanswered. First, the proteome of the administered cells was not characterized, and functional studies to identify the specific drivers of the observed recovery were not conducted. As a result, the precise mechanisms underlying the therapeutic effects remain undefined, limiting the ability to target these pathways for optimization. Second, this study focused on the glial cell response, which is only one aspect of the complex pathophysiology of HIE. Other key contributors, such as neuronal remodeling, brain connectivity, vascular remodeling, and immune system interactions, should be further explored to enhance our understanding of the multifaceted recovery process. Third, the single time point of proteomic analysis, 30 days post-injury (i.e. 28 days post-cell administration), restricts observations to the long-term effects of the treatment. While this is informative for understanding sustained recovery mechanisms, it does not allow for the detection of direct cellular actions in the acute or subacute phases after administration. Some of the observed changes may be secondary effects, driven by the improved overall brain status rather than the cells’ direct influence. Nonetheless, this work establishes a robust foundation for advancing stem cell-based therapies in neonatal HIE, with direct implications for clinical translation.

## 5. Methods

### 5.1. Ethical Approval

Approval for animal studies was obtained from the Ethical Committee of the University of Beira Interior and authorized by the Portuguese General Directorate for Food and Veterinary (0421/000/000/2019). The research adhered to the European Directive (2010/63/EU) governing the protection of laboratory animals used for scientific purposes. Additionally, ethical approval for using human mesenchymal stem cells was granted by the Ethical Committee of the Faculty of Medicine of the University of Coimbra (Approval Number: 075-CE-2019). The use of animals in this project is justified by the importance of assessing the impact of the proposed strategies on functional outcomes, such as cognitive and motor capabilities, for their translation to clinical applications. The animal research conducted in this study is reported following the Animal Research: Reporting of In Vivo Experiments (ARRIVE) 2.0 guidelines (73).

### 5.2. Animals

Animal studies were carried out in the animal facility of the Faculty of Health Sciences, University of Beira Interior (Covilhã, Portugal). Animals enrolled in this study were maintained in an alternating 12-hour light/dark cycle, maintained with their dam until weaning at P21, and handled daily after the induction of the neonatal HI injury to monitor welfare. Experimental design, including sample size calculation, was done using Experimental Design Assistant (https://eda.nc3rs.org.uk). The effect size and variability were determined by previous experiments from the group. The individual animal served as the experimental unit and was randomly and independently assigned to one of the experimental groups with sex as a blocking factor. The researchers were not blinded for the behavioral tests or statistical analysis.

### 5.3. Induction of neonatal HI brain injury

Male and female Wistar rat pups at P10 (weighting 15-20 g) were subjected to an HI brain lesion following the adapted Rice-Vannucci model for HIE. First, the animals were anesthetized using isoflurane (5% for induction, 1.5-2% for maintenance; Isoflo, Zoetis). Them, the left common carotid artery was exposed and ligated using 6−0 silk suture (F.S.T).

For the hypoxia, the animals were placed in an airtight chamber at 37°C filled with a mixture of 8% oxygen and 92% nitrogen (Air Liquide) for 90 minutes. Control animals underwent a similar procedure; however, their left common carotid artery was not ligated and were exposed to room air in a heated chamber at 37°C.

### 5.4. UC-MSCs culture

UC-MSCs were isolated from the Wharton’s Jelly of cryopreserved fragments of human UC, as previously described (74). When colonies were observed, cells were washed with PBS, detached by the addition of 0.05% Trypsin-EDTA (Gibco™) on a humidified incubator at 37°C. Then, UC-MSCs were homogenized and centrifuged at 290×g, at room temperature. The cell pellet was resuspended and UC-MSCs were replated in Minimum essential Medium-α (Gibco™) supplemented with 5% (v/v) fibrinogen depleted human platelet lysate (HPL) (UltraGRO™, Helios) and Antibiotic-Antimycotic (Gibco™), until sub confluence was achieved. This procedure was repeated until passage four and UC-MSCs were cryopreserved in HPL with 10% dimethyl sulfoxide.

### 5.5. Hypoxic preconditioning of UC-MSCs

UC-MSCs were plated in Minimum Essential Medium-α (Gibco™) supplemented with 5% (v/v) fibrinogen depleted HPL (UltraGRO™, Helios) and Antibiotic-Antimycotic (Gibco™), until sub confluence was achieved. This procedure was repeated to expand the number of cells in culture. For hypoxic preconditioning, UC-MSCs were kept at standard culture conditions until passage three. Then, 24 hours before the stimuli, cells were seeded at 10,000/cm^2^ (i.e. passage four). Before the stimuli, the culture media was washed with PBS and replaced with Minimum Essential Medium-α (Gibco™) without supplementation, and UC-MSCs were placed for 24 hours on a humidified incubator (Binder), at 37°C, with 5% CO2 (N-MSCs) – or in an InvivO₂® 400 humidified incubator (Baker Ruskinn), set at 0.1% O_2_/5% CO_2_ (SSH-MSCs) or 5% O_2_/5% CO_2_ (MH-MSCs). Preconditioned UC-MSCs were cryopreserved in HPL with 10% dimethyl sulfoxide and stored in liquid nitrogen until administration.

### 5.6. UC-MSCs administration

MSCs were delivered by IV route two days after the induction of HI brain lesion (i.e. P12). On the day of administration, the previously prepared MSCs were thawed and centrifuged at 290×g for 5 minutes. After removing the supernatant, MSCs were resuspended in PBS and viable cellular density was determined using the Trypan Blue exclusion method.

For IV administration, the animals were anesthetized with isoflurane and viable UC-MSCs diluted in 200 µl of PBS were administered, per animal, in the tail vein using a 29-gauge insulin syringe (Terumo). Each animal receiving UC-MSCs was administered with 50,000 cells.

### 5.7. Behavioral analysis

#### 5.7.1. Negative Geotaxis Reflex

The negative geotaxis reflex test was used to assess rat’s motor coordination early in development, at P14 and P17. For this test, rat pups were placed downhill on a 45° slanted slope, and the time required for the pups to face uphill was recorded. No animals were excluded from the analysis.

#### 5.7.2. Novel Object Recognition Test

The recognition memory of the animals was assessed using the novel object recognition test at two developmental stages: infancy (P21) and early adulthood (P38) (75). Prior to the test day, animals underwent a habituation phase in the testing arena, lasting 10 minutes. On the test day, animals were initially presented with two identical objects during a 10-minute familiarization phase. After a 30-minute interval, the animals were exposed to one familiar object and one novel object for 10 minutes during the test phase. Exploration times for both the familiar and novel objects were recorded during the initial five minutes of the test phase. Subsequently, a discrimination ratio was computed using the formula: (time spent exploring the novel object) / (total exploration time) × 100. A similar procedure was followed for the assessment at P38, except that different sets of familiar and novel objects were used.

#### 5.7.3. Footprint test

As previously described, the footprint test served as the primary method for identifying locomotor deficits and gait abnormalities (72, 76). The assessment was conducted at P28, corresponding to a developmental stage akin to human childhood (75). Rats’ fore and hind paws were coated with non-toxic paint, following which they were prompted to cross a 100 cm path lined with paper, leading towards a black box. Evaluation of the footprint pattern entailed analyzing ten consecutive steps, during which instances of foot-dragging or overlapping footprints of the contralesional paws were counted. Animals showing a lack of motivation to cross the corridor were excluded from subsequent analysis.

#### 5.7.4. Ladder Rung Walking Test

The assessment of animals’ motor function, particularly coordination, was conducted using the ladder rung walking test at P30, a developmental stage corresponding to human childhood (75). The apparatus for this test consisted of a 100 cm long corridor delineated by two transparent acrylic side walls, between which metal rods (0.3 cm in diameter) were positioned 1 cm apart. Rats crossed the apparatus four consecutive times under video recording. Subsequently, each video was reviewed in slow motion to count the number of foot slips occurring between the rods. Animals showing a lack of motivation to cross the ladder were excluded from subsequent analysis.

### 5.8. Tissue Collection and Preparation

At postnatal day 40, the animals were euthanized with an overdose of anesthetics (200 mg/kg Ketamine and 10 mg/kg Xylazine), followed by perfusion with PBS and 4% paraformaldehyde. Following perfusion, the brains were excised and immersed in 4% paraformaldehyde for 16 hours at 4 °C. Subsequently, for cryopreservation purposes, the brains were transferred to a 30% sucrose solution until they sank. Once cryopreserved, the brain tissue was rapidly frozen in liquid nitrogen and stored at −80 °C until further processing. Sectioning of the frozen brains was performed using a cryostat (Leica CM3050) set to a thickness of 40 µm, with sections collected at 240 µm intervals.

### 5.9. Cresyl Violet Staining

Brain lesion extension was assessed in sixteen sequential brain sections for each animal. Frozen brain sections were mounted in poly-lysin coated glass slides (VWR) and stained with 0.05% Cresyl Violet Acetate (Merck) using the Sakura TissueTek Automated DRS 2000 automated slide stainer. Images were acquired with the Axio Imager A1 Microscope (Zeiss) with a 5× objective (EC Plan-Neofluar 4×/0.16 M27), and the volume of the left (ipsilesional) and right (contralesional) hemispheres were determined with the Cavalieri’s Principle Probe of the StereoInvestigator software (MBF Bioscience). Brain lesion extension was calculated as (V_contralesional_ − V_ipsilesional_)/V_contralesional_ × 100.

### 5.10. Proteomic analysis of brain tissue

#### 5.10.1. Brain tissue collection

At P40, the animals were anesthetized with isoflurane (Isoflo, Zoetis), followed by cervical dislocation and decapitation. Ipsilesional and contralesional brain tissue of the rats was frozen in liquid nitrogen, and stored at -80 °C. For each experimental condition, eight samples were collected, each corresponding to an individual animal, except the control group which had six samples.

#### 5.10.2. Sample preparation – ShortGeLC and Peptide Extraction

##### Homogenization of brain tissue samples

Upon thawing, the cerebellum and brainstem were removed from each sample. Each tissue sample was placed in a microcentrifuge tube and weighed. Tissue samples were homogenized in 2× Laemmli Sample Buffer (SB; 0.12 M Tris-HCl pH 6.8, 3.33% (w/v) sodium dodecyl sulfate, 10% (v/v) glycerol, 3.1% (w/v) dithiothreitol, and bromophenol blue) at a ratio of 2.5 ml buffer per 600 mg of tissue. Samples were sonicated twice for 40 seconds each (3 seconds ON, 2 seconds OFF) at 60% amplitude. Between samples, the sonicator probe was cleaned with ddH2O and 96% ethanol. The homogenates were boiled at 95°C for 10 minutes with agitation at 300 rpm. After denaturation, samples were alkylated with 40% acrylamide (Biorad). The samples were centrifuged at 15,000 × g for 15 minutes. Samples were immediately used for protein quantification and remaining steps or stored at -80 °C.

##### Protein quantification

Protein concentration was determined using the Pierce™ 660nm Protein Assay Kit (Thermo Fisher Scientific) with Ionic Detergent Compatibility Reagent (Thermo Fisher Scientific). The assay reagent was prepared by adding 1 g of Ionic Detergent Compatibility Reagent per 20 ml of Pierce reagent. In a 96-well microplate, 10 μl of either 2× SB (for standard curve) or water (for samples) was combined with 10 μl of protein standard or diluted sample (1:20 dilution). The reaction was initiated by adding 150 μl of prepared reagent. The plate was sealed and shaken at 500 rpm before measuring absorbance at 660 nm. A calibration curve was generated using six protein standards, and this curve was used to determine the protein concentration in each experimental sample. Two micrograms of the recombinant protein Green Fluorescent Protein (GFP) was added to each sample as an internal standard.

##### Short GeLC-MS - in gel protein digestion protocol

Protein samples were separated using a partially electrophoretic separation protocol as previously described – Short GeLC-MS (77). The running buffer was prepared by diluting 100 mL of 10× Tris/Glycine/SDS buffer (Bio-Rad) in 900 mL of double-distilled water. The Mini-PROTEAN® Tetra cell system (Bio-rad) was assembled according to the manufacturer’s instructions. The electrophoresis chamber was filled with 1× running buffer to the indicated mark, and polyacrylamide gel (4-20% TGX Stain-Free Gel, Bio-rad) wells were rinsed with running buffer using a pipette. Protein samples (50 μg total protein of each sample for quantification) or pools of each experimental condition (70 μg total protein of each pool for identification) and molecular weight markers were loaded into the wells, with any empty wells being filled with an equal volume of 2× Laemmli buffer. Electrophoresis was performed at 110 V for approximately 18 minutes. After electrophoresis, proteins were visualized with Colloidal Coomassie Blue (78). Briefly, the gel was rinsed in distilled water and submerged in a staining solution (10% (v/v) of 85% phosphoric acid, 10% (w/v) ammonium sulfate, 20% (v/v) methanol and 0.2% Coomassie Brilliant Blue G-250 (Thermo Fisher Scientific) for approximately 1 hour under orbital rocking platform. After staining, gels were transferred to fresh, pre-cleaned containers and washed repeatedly with distilled water until the background was clear. Final destaining was performed overnight on a rocking platform with distilled water. Gel processing was done the following day. To maintain gel integrity and prevent contamination, gloves were washed with SDS detergent throughout the procedure as needed, acetate sheets were cleaned sequentially with detergent, water, and ethanol, and band excision was done in clean acetate sheets placed in a laminar flow chamber. Each lane was sliced into bands of equal size. Following excision, each band was sliced into small pieces and transferred to 96-well deep-well plates with 600 µL of distilled water.

Gel pieces in 96-well deep-well plates were treated with 600 µL of destaining solution (50 mM ammonium bicarbonate and 30% acetonitrile). Samples were incubated under agitation at 700 rpm for 15 minutes. The destaining solution was removed, and the process was repeated until the gel pieces appeared transparent. Following destaining, gel pieces were washed once with an equal volume of water with agitation. Then, gel pieces were dried in a SpeedVac concentrator at 60°C for 1 hour. After dehydration, 0.01 mg/mL trypsin in 10 mM ammonium bicarbonate was added to each well and plate was incubated for 15 minutes. After this initial incubation, more 10 mM ammonium bicarbonate buffer was added to the wells and trypsin was let to react overnight at room temperature in the dark (approximately 16 hours).

Following enzymatic digestion, peptides were extracted using a sequential extraction protocol with increasing acetonitrile concentrations. Three extraction solutions were prepared: Solution A (30% acetonitrile with 1% formic acid), Solution B (50% acetonitrile with 1% formic acid), and Solution C (98% acetonitrile with 1% formic acid). The initial digestion supernatant containing released peptides was transferred to a 1.5 mL low-binding microcentrifuge tube. Sequential extractions were then performed by adding 100 µL of each extraction solution (A, B, and C) to the gel pieces, with 15-minute incubations at 25°C and 700 rpm agitation in a thermomixer for each step. After each extraction, the supernatant was collected and combined with the initial digest solution in the same microcentrifuge tube. For data-independent acquisition (DIA)-based sequential window acquisition of all theoretical mass spectra (SWATH-MS) analysis or SWATH/DIA-MS, peptides from all gel sections were pooled for each sample, while for data-dependent acquisition mode (DDA) and library generation, the gel sections were processed and maintained separately. Following the completion of all extraction steps, the combined extracts were dried in a SpeedVac concentrator at 60°C until complete dryness was achieved.

#### 5.10.3. Mass spectrometry

Dried peptide samples were reconstituted in a solution containing 2% acetonitrile and 0.1% formic acid before analysis on NanoLC™ 425 System (Eksigent) coupled to a Triple TOF™ 6600 mass spectrometer (Sciex) with an electrospray ionization source (DuoSpray™ Ion Source from Sciex). All data were acquired using Analyst TF 1.8.1 software (Sciex).

Chromatographic separation of peptides was carried out using a Triart C18 Capillary Column 1/32” (12 nm, 3 μm, 150 mm × 0.3 mm, YMC, Dinslaken, Germany) and a Triart C18 Capillary Guard Column (0.5 mm × 5 mm, 3 μm, 12 nm, YMC) at 50 °C. The flow rate for the separation process was set at 5 µL/min. The mobile phases used for the gradient were mobile phase A (0.1% formic acid and 5% dimethyl sulfoxide in water) and mobile phase B (0.1% formic acid and 5% dimethyl sulfoxide in acetonitrile). The separation was performed with a gradient elution program, which involved a gradual increase in mobile phase B from 5% to 30% over the first fifty minutes, followed by a rapid ramp from 30% to 98% over two minutes (50-52 min), 98% over two minutes (52-54 minutes), 98% to 5% over two minutes (54-56 minutes), and equilibration phase during 10 minutes with mobile phase B with 5% acetonitrile. The ESI DuoSpray™ ionization source was operated in positive ion mode. Key instrument parameters included an ion spray voltage of 5500 V, and both the nebulizer gas (GS1) and curtain gas (CUR) were maintained at 25 psi.

In DDA mode, full MS spectra were collected across a mass range of m/z 350-2250 with an accumulation time of 250 ms, followed by up to 100 MS/MS scans (m/z 100-1500) with a 30 ms accumulation time. The total cycle time was maintained at 3.3 s. Precursor ion selection was based on charge states between +1 and +5, with a minimum intensity threshold of 10 counts per second. In SWATH/DIA-MS mode, the mass spectrometer was operated in a looped production mode and specifically tuned to a set of 168 overlapping windows, covering the precursor mass range of 350-1250 m/z. A 50 ms survey scan (350-1250 m/z) is acquired at the beginning of each cycle, and SWATH-MS/MS spectra were collected from 100-1500 m/z for 20 ms resulting in a cycle time of 3.3 seconds.

#### 5.10.4. Proteomics Data Analysis

Each band of pooled samples was acquired individually in DDA mode. The ion library of the precursor masses and fragment ions was generated in ProteinPilot^TM^ (Sciex) by combining all files from the experimental group pools, and data was searched using the reviewed Rattus norvegicus Swiss-Prot database. An independent False Discovery Rate (FDR) analysis was used to assess the quality of identifications, with a target-decoy approach.

PeakView^TM^ (Sciex) was used for the analysis of data obtained with SWATH/DIA-MS acquisition and to retrieve the relative protein quantification. First, the library generated with the DDA data was imported into the software, indicating the number of proteins obtained with a local FDR below 5%. Then, the SWATH/DIA-MS data files corresponding to each sample were loaded and processing was done with the unique peptides and an FDR below 1%. Data was normalized to the total intensity.

Statistical analysis of the proteomics data was performed on MetaboAnalyst 6.0 (79) and RStudio 2024.09.0. Proteins of interest were selected through multivariate partial least squares-discriminant analysis (PLS-DA) and univariate Kruskal-Wallis test analysis. Differentially expressed proteins (DEPs) had a variable important in projection (VIP) score > 1 (in component 1 or 2) and p-value < 0.05. DEPs were explored with Gene Ontology Enrichment analysis in PANTHER Classification System 19.0 (80–82), Reactome Pathway Knowledgebase (83) , and protein-protein interactions (PPI) were identified with STRING database 12.0 (84). Gene Ontologies were visualized with ImageGP (85), Venn diagrams with BioVenn (86), and violin plots with GraphPad Prism 10 (GraphPad Software Inc., San Diego, CA).

### 5.11. Statistical Analysis (except proteomics data)

Statistical analysis was performed using GraphPad Prism 10 (GraphPad Software Inc., San Diego, CA). Outliers from each data set were identified with the Grubbs’ analysis (α=0.05) and excluded from the analysis. Statistical analysis was performed with the Kruskal Wallis test coupled with Dunn’s multiple comparison correction test or One-way ANOVA coupled with Tukey’s multiple comparison test, according to the type of data. The differences were considered significant when the p-value was inferior to 0.05. Details of the statistical analysis can be found in the figure legends.

## Notes

### Competing Interest Statement

The authors have declared no competing interest.

