## Supplementary Figure 1 for "Quieting the Storm: Hypoxia as a Strategy to Boost UC-MSC Therapies for Neonatal Hypoxic-Ischemic Encephalopathy"

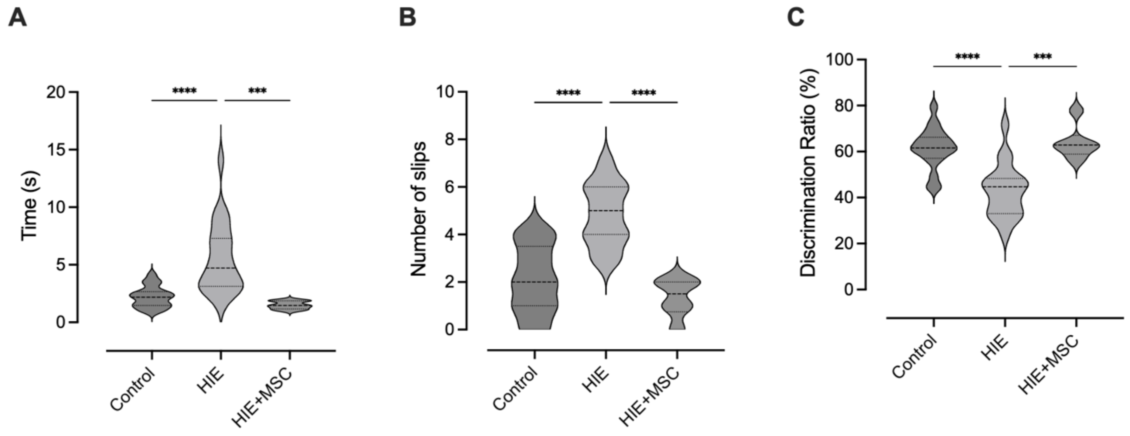


**Supplementary Figure 1.** Intravenous administration of 0.5 x $10^6$ MSCs rescues motor and cognitive impairments after neonatal hypoxic-ischemic brain injury.
(A) The negative geotaxis reflex was performed at postnatal day 17 (P17) to evaluate sensorimotor function.
(B) The ladder ung walking test was used to evaluate motor coordination and the number of slips that the animals had during the test was accounted for at P30.
(C) The novel object recognition test was employed to evaluate memory deficits at P38.
Data is presented as violin plots and statistical analysis was performed using one-way ANOVA with Tukey's multiple comparison correction (*** $p < 0.001$, **** $p < 0.0001$).
